# Epidemiological and space aspects of the schools of the National Leprosy Campaign: the case of Sobral – Ceará, Brazil

**DOI:** 10.1101/663229

**Authors:** Vitória Ferreira do Amaral, Maria Socorro Carneiro Linhares, Francisco Rosemiro Guimarães Ximenes Neto, Sandra Maria Carneiro Flor, Ligia Regina Franco Sansigolo Kerr, Luíza Jocymara Lima Freire Dias, Isabel Cristina Kowal Olm Cunha, Neyson Pinheiro Freire, Manoel Carlos Neri da Silva, Marcos Aguiar Ribeiro, Izabelle Mont’Alverne Napoleão Albuquerque, Ana Suelen Pedroza Cavalcante

**Author notes:** Corresponding author (VFA). 1. Conceptualization: Original Draft Preparation, Data Curation, Formal Analysis, Review & Editing, Investigation, Methodology, Resources, Validation, Visualization. 2. Conceptualization: Data Curation, Formal Analysis, Review & Editing, Investigation, Methodology, Project Administration, Resources, Supervision, Validation. 3. Conceptualization: Data Curation, Formal Analysis, Review & Editing, Investigation, Methodology, Project Administration, Resources, Supervision, Validation. 4. Conceptualization: Data Curation, Formal Analysis, Review & Editing, Investigation, Methodology, Project Administration, Resources, Supervision, Validation. 5. Review & Editing: Resources, Formal Analysis. 6. Review & Editing 7. Review & Editing: Resources, Formal Analysis, Review & Editing. 8. Review & Editing 9. Review & Editing 10. Review & Editing: Resources, Formal Analysis, Review & Editing. 11. Review & Editing: Resources, Formal Analysis, Review & Editing. 12. Review & Editing.

## Abstract

The objective of this study is to describe the epidemiological and spatial aspects of leprosy of the schoolchildren participating in the National Leprosy Campaign in the municipality of Sobral, Ceará, in the year 2016. This is a cross-sectional epidemiological study with spatial analysis of secondary data obtained in the records instruments used in the Campaign (self-image files), with public school schoolchildren from five to fourteen years old. From this population a sample was taken to be studied from the calculation of a standard error limit of 5%, confidence level of 99.99% and an expected frequency of 50%, resulting in 1,216 students, corresponding to 19.7% of a total of 6,169 schoolchildren who returned the completed self-indexed records. A descriptive analysis was performed for all the variables of interest of the study object and for the spatial analysis the QGIS program 2.18 was used. Of the 1,216 schoolchildren participating in the study, 31.7% had body spots and 18.1% (220/1126) of the total number of schoolchildren had leprosy cases in the study. Of the 1,216 students in the study, 31.7% (386/1). Among the schoolchildren with spots on the body, 6.2% (75/1126) reported having cases of leprosy in the family, 59.3% (195/329) are birthmarks, 20.7% Among children with spots suspected of leprosy (39.2%, 129/329), they were found to be dormant (9.3%) (31/329) and 10.6% (35/329) The strategies for the screening of new suspected leprosy cases developed through campaigns helped to mobilize around the epidemiological situation of leprosy, facilitating the dissemination of information to leprosy patients. the population on the recognition of signs and symptoms, treatment and cure of leprosy.

**AUTHOR SUMMARY:** Leprosy is an infectious and contagious, chronic disease caused by Mycobacterium leprae (M. leprae), which has high infectivity and low pathogenicity. Brazil is part of the group of three countries responsible for 80.2% of all new cases registered in the world in 2017 and in the Region of the Americas contributed with 92.3% of new cases. The National Leprosy Campaign aims to find new cases of leprosy in children 5 (five) to 14 years of age. Sobral, a city of Ceará, with high disease burden, has been joining the campaign every year since 2013. This study describes the epidemiological and spatial aspects of leprosy of students participating in the National Leprosy Campaign in the municipality of Sobral, Ceará, in 2016. Strategies for screening new suspected leprosy cases developed through Campaigns, in addition to contributing to the identification of new cases in the community, promote a mobilization around the epidemiological disease situation and dissemination of information to the population on the recognition of signs and symptoms, treatment and cure of leprosy.

## INTRODUCTION

Leprosy is an infectious and contagious, chronic disease caused by Mycobacterium leprae (M. leprae), which has high infectivity and low pathogenicity. Transmission in humans occurs mainly through the upper airways with the elimination of the M. leprae agent, which is preferentially installed in the skin and peripheral nerves ^1^. Its multiplication in the organism is slow, which determines a long period of incubation of the disease, which can vary from two to seven years ^2^, and the later the disease is diagnosed, the greater the risk of developing deformities and incapacities ^3^.

The number of new cases of leprosy declines worldwide, but in 2017, in Brazil there was an increase in notifications, with 1,657 new cases. In the Region of the Americas, Brazil contributed 92.3% of new cases in the last year. Only Brazil, India and Indonesia represent 80.2% of all registered cases worldwide. In Brazil, children represented 8.06% of the total number of new cases reported, which may mean failure to detect cases in adults and persistence in the disease transmission cycle ^4^.

As a measure to strengthen the fight against leprosy, since 1992, the World Health Organization (WHO) has been launching campaigns to eliminate the disease as a public health problem, as control strategies where the burden of disease is high. In 2016, the “Global Strategy for Leprosy: 2016-2020” was launched to accelerate action towards a world without leprosy, aiming at strategic changes in three areas: (1) elimination of Grade 2 disabilities in children with leprosy; (2) reduction of new cases of leprosy with Grade 2 disabilities to less than one case per million population; (3) that no country has laws that allow discrimination to those with leprosy ^5,6^.

In Brazil, new strategies and intervention measures were introduced to achieve the elimination of neglected diseases by establishing the “Integrated Plan for Strategic Actions to Eliminate Leprosy, Filariasis, Schistosomiasis and Onchocerciasis as a Public Health Problem.” In this plan, the active search for new cases of leprosy disease from the self-image files for leprosy, applied to schoolchildren ranging from five to fourteen years old, 7 from campaigns in public schools, which are characterized as interdisciplinary spaces of construction of knowledge, and that can enhance the insertion of new practices and habits, constituting the appropriate tool for the success of educational actions in health ^8,9^.

The National Leprosy, Verminoses, Trachoma and Schistosomiasis Campaign has annual periocity and focuses on geographic areas with greater epidemiological risks for leprosy, with the search for new cases in children between five and 14 years old, aiming at the early detection through use of the “mirror method”, which consists in the completion of a self-image record by parents or guardians indicating whether or not the child has suspicious spots of leprosy ^8,10^.

Sobral, one of the priority municipalities in the state of Ceará for surveillance and leprosy control actions since 2013, has been joining the annual campaigns of Leprosy, Verminoses, Trachoma and Schistosomiasis of the Ministry of Health as a strategy for the early detection of new cases of leprosy, and to minimize the endemic load. This municipality, historically, presents high annual rates of detection of new cases of leprosy. Between 2005 and 2016, 1,163 new cases were registered in this municipality, with an average detection rate of 52.2 cases per 100,000 inhabitants. Of the total number of cases identified in the period, 89 were younger than 15 years, reaching an average detection rate of 10.2 cases per 100,000 inhabitants in this age group ^11^. Mean values in the general population and in those under 15 indicate, according to the parameters of the Ministry of Health, as a hyperendemic municipality ^10^.

This study aims to describe the epidemiological aspects of the students of the public school of Sobral, aged between five and 14 years old, who participated in the National Leprosy Campaign in 2016.

## METHODOLOGY

### Ethics statement

This study is part of a larger study entitled: Leprosy in children under 15 years: use of social networks and genetic tools in the study of the transmission of Mycobacterium leprae, which was approved by the Scientific Committee of the Secretariat of Health and Social Action of Sobral and by the Human Research Ethics Committee of the Federal University of Ceará (CEPSH-UFC), Nº. 624,393.

During the National Leprosy Campaign the parents received a consciously printed informational statement accompanying the self-image file for leprosy. In the end it is explained that the objective of the campaign is to protect children from diseases: leprosy, verminoses, trachoma and schistosomiasis. Parents who accepted, their respective children were evaluated for identification of suspected cases of leprosy.

### National Leprosy Campaign

This is a cross-sectional epidemiological study with spatial analysis of secondary data obtained in the National Campaign on Leprosy, Verminoses, Trachoma and Schistosomiasis, with students from the public network of the municipality of Sobral, in the state of Ceará-Brazil, in 2016 The epidemiological scenario of leprosy in Sobral was one of the motivators for choosing the place for the present case study.

Epidemiological “cross-sectional” studies seek to give an idea of the cut in the historical flow of a disease, evidencing its characteristics and correlations at that moment, in a given population or community^12^. While the spatial analysis based on the use of the Geographic Information System (SIG) in epidemiology allows the analysis of the spatial distribution, predicting patterns, determinants and results directed to populations ^13^. Allowing the understanding of the epidemiological dynamics of the territory from the design of a cartographic chart.

The population of the study are schoolchildren in the age range from five to fourteen years of the 31 schools in the municipal network, of which 22 are urban and nine are in the countryside. The schools that joined the campaign were selected based on two criteria: to include a higher concentration of students enrolled in the study’s age range, and to be located in areas with high rates of leprosy cases.

The operation of the campaign in the schools comprised four stages: 1st Step – identification of the Family Health Centers (CSF) of reference of the schools; Step 2 – CSF and school health professionals received guidelines and campaign materials, such as self-image files for leprosy; Step 3 – delivery of self-image files to schoolchildren and completion by the parents or persons in charge at home; Step 4 – return of the self-image files to the health professionals of the responsible schools within a maximum of two days, and screening of schoolchildren with signs and symptoms suggestive of leprosy who need to be referred for medical referral in the school’s reference CSF.

In the campaign, 19,415 self-image files were distributed to schoolchildren, and 6,169 of the files were returned by the schoolchildren. Considering the significant quantity of returned chips, the sample was calculated in the public domain software Epi Info ™, with a default error limit of 5%, a 99.99% confidence level and an expected frequency of 50%, resulting in 1,216 tokens of self-image selected by the random sampling technique, corresponding to 19.7% of the total of filled in received.

In order to access and use the data of the self-image files of the students of the National Leprosy Campaign in 2016, the Health Surveillance Coordination of the municipality of Sobral, local responsible for the Campaign, was requested by means of a Term of Commitment. Thus, as a criterion for the selection of self-image files, access was made to the self-image files returned and the correct completion. The records that showed inconsistencies were not selected for analysis.

The analysis variables from the self-image dataset were the sociodemographic data of the schoolchildren and the investigative questions of the characteristics of the spot, such as: If there is a spot on the skin? If the spot is from birth? If the spot itches? If the spot hurts? If the spot is dormant? And the family history of leprosy.

From the variables, groups were stratified to analyze the characteristics of the spots, being defined as: suspicious spot – schoolchildren with spots on the skin that are not of birth, itches and do not hurt; suspect spotting and case of leprosy in the family – schoolchildren with spots on the skin that are not birthmarks, do not itch and do not hurts and have cases of leprosy in the family; spotting dormant suspect-schoolchildren with dormant spots that are not birthmarks; suspicious spot with cases of leprosy in the family – schoolchildren with dormant spots that are not birthmarks and cases of leprosy in the family.

In the spatial analysis, maps of the main characteristics of the schoolchildren correlated to the adult leprosy rate of 2016 in Sobral were constructed using the Voronoi diagram and the linear and proximity distance matrix. The Voronoi or Polygon diagrams of Theissen are categories of maps that produce spatial mosaic around a set of points, of a space, that are characterized as areas of influence of a point in the space occupied in space mosaic ^14^. As method of analysis, was used the vector of the linear distance matrix (N * k × 3), the closest neighbor calculation and the polygonal point count were applied.

For the construction of the map, the cartographic base of the Brazilian Institute of Geography and Statistics (IBGE) was used from the census shapefiles of the sectors. The information geoprocessing and map construction were carried out in the public domain program QGIS 2.18. Schoolchildren residing in the urban area were georeferenced by online software MyGeoPosition (http://my.gageoposition.com/) where they set latitude and longitude of each address for positioning on maps. While the coutryside schoolchildren were not mapped, because the geocodes of the residential addresses were not available, and it was not possible to georeference.

Data for the calculation of leprosy detection rates for the year 2016 were collected in the municipal database of the Notification of Injury Information System (SINAN) and the 2010 population census by sector of IBGE (2010). As a measure of establishing more precise data for the calculation of the leprosy rate for the year 2016 was applied the calculation of population intercensal projection by IBGE.

The detection rates of Sobral neighborhoods were presented on maps based on the endemicity classification parameters of the Ministry of Health’s Leprosy Elimination Progress Monitoring Indicators, which establishes a new case detection coefficient ≥ 40.00 / 100,000 inhabitants as hyperendemic; very high: 20.00 to 39.99 / 100,000 inhabitants; high: 10.00 to 19.99 / 100,000 inhabitants; average: 2.00 to 9.99 / 100,000 inhabitants. and low: <2.0 / 100,000 inhabitants ^15^.

Statistical analysis of the variables was performed in the Statistical Package for Social Sciences (SPSS) software from a set of tools for data analysis using descriptive statistical techniques and the Pearson’s Chi-square test with a level of significance of 0.05% and confidence of 95% for sociodemographic variables and related to leprosy research in the self – image record.

## RESULTS

Of the 1,216 schoolchildren in the study, 31.7% (386/1126) had spots on the body and 18.1% (220/1126) of the total number of schoolchildren had leprosy cases in the family. Among students with spots on the body, 6.2% (75 / 1,216) reported having cases of leprosy in the family. As for the sociodemographic characteristics, the majority of schoolchildren in the study are female, 56.4% (686 / 1.216), and 52.3% (636 / 1.216) present a prevalent mean age of 9.4 years (standard deviation [58]), with 56.6% (689 / 1.216) attending Elementary School I (Table 1).

**Table 1.** Socio-demographic characteristics of the students of the IV National Leprosy Campaign in Sobral – Ceará, Brazil, 2018.

| Categories / Variables | schoolchildren | schoolchildren with spots on the body | schoolchildren with cases of leprosy in the family | schoolchildren with spots on the body and cases of leprosy in the family | p-value |
| --- | --- | --- | --- | --- | --- |
| (n, %) | n=1.216 (%) | n=386 (31,7%) | n=220 (18,1%) | n=75 (6,2%) |  |
| Gender |  |  |  |  |  |
| Female | 686 (56,4) | 167 (43,3) | 116 (52,7) | 36 (48,0) | 0,129 |
| Male | 530 (43,6) | 219 (56,7) | 104 (47,3) | 39 (52,0) |  |
| Age group |  |  |  |  |  |
| 5 to 9 years | 636 (52,3) | 192 (49,7) | 124 (56,4) | 41 (54,7) | 0,672 |
| 10 to 15 years | 580 (47,7) | 194 (50,3) | 96 (43,6) | 52 (69,3) |  |
| ESchooling |  |  |  |  |  |
| Children's Ed | 88 (7,2) | 26 (6,7) | 15 (6,8) | 02 (2,7) | 0,270 |
| Elementary school | 689 (56,6) | 210 (54,4) | 135 (61,4) | 46 (61,3) |  |
| Middle school | 439 (36,1) | 150 (39) | 70 (31,8) | 27 (36,0) |  |
| Localization |  |  |  |  |  |
| Urban area | 909 (74,8) | 317 (82) | 178 (80,9) | 70 (93,3) | 0,001 |
| Countryside | 307 (25,2) | 69 (17,9) | 42 (19,1) | 05 (6,7) |  |

\* Pearson's Chi-square test of schoolchildren with blemishes and cases of leprosy in the family.

Among the schoolchildren who reported on the self-image card, 56.7% (219/386) were male, but this result was not statistically significant between sex proportions (p = 0.129), of that group 50.3% (194/386) are in the age range of 10 to 15 years, and 54.4% (210/386) studied in elementary school-I (Table 1).

While schoolchildren with cases of leprosy in the family, 52.7% (116/220) are female, 56.4 (124/220) are between five and nine years of age, 61.4% (135/220) attended the Elementary School I and 80.9% (178/220) lived in the urban zone (Table 6). Of the group of schoolchildren with a lesion in the body, 52% (39/75) of cases of leprosy are male, 69.3% (52/75) are in the age range of 10 to 15 years, 61.3 % (46/75) attend Elementary School I (Table 1).

The analysis of the location of the students’ residence had significant statistical significance (p <0.001), with 74.8% (909 / 1,216) of schoolchildren living in the urban area, and 93.3% (70/77) had spots and cases of leprosy in the family (Table 1).

With the distribution of the types of spots, the sociodemographic characteristics of the schoolchildren with birthmarks, spots that itches, spots that hurt and spots that are dormant of the total of 329 schoolchildren who indicated in the self-image record some of these defining characteristics were analyzed. (Table2). It is worth noting that 14.7% (57/386) of schoolchildren indicated that they have a spot on the body, they are not: birthmarks, do not itch, do not ache and are not dormant.

Among schoolchildren who presented spots with a defining characteristic, 59.3% (195/329) were birthmarks, 20.7% (68/329) itch, 9.4% (31/329), and 10.6 % (35/329) are dormant (Table 2). Of the schoolchildren with birthmarks, 58% (113/195) were males, of which 51.3% (100/195) were in the age range of 10 to 15 years, 52.3% (102/195) studied in the Elementary School I and 76.4% (149/195) lived in the urban area. Schoolchildren with spots that itch, 60.3 (41/68) are males, 56% (38/68) are between 5 (five) and 9 years old, 58.8 (40/68) enrolled in elementary school I and 84% (57/68) lived in the urban area. And schoolchildren with spots that aches, 61.3% (19/31) are males, 68% (21/31) were in the age range of 5 (five) to 9 years, 80.7% (25/31) enrolled in Elementary School I and 93.5% (29/31) of schoolchildren lived in the urban zone (Table 2).

**Table 2.** Distribution of the characteristics of the spots of the students of the IV National Leprosy Campaign according to sex, age group, level of schooling and location, in the municipality of Sobral – Ceará, Brazil, 2018.

| <u>Spots*</u> |  |  |  |  |  |  |
| --- | --- | --- | --- | --- | --- | --- |
| Categories / Variables | Birthmark | spot itches | spot hurts | dormant spot | spot without characterization | **<br>p-value |
| (n, %) | n=195 (59,3%) | n=68 (20,7%) | n=31 (9,4%) | n=35 (10,6%) | n=57 (14,7) |  |
| **Gender** |  |  |  |  |  |  |
| Female | 82 (42,0) | 27 (39,7) | 12 (38,7) | 17 (48,6) | 29 (50,9) | 0,546 |
| Male | 113 (58,0) | 41 (60,3) | 19 (61,3) | 18 (51,4) | 28 (49,1) |  |
| **Age Group** |  |  |  |  |  |  |
| 5 to 9 years | 95 (48,7) | 38 (56,0) | 21 (68,0) | 13 (37,1) | 25 (43,9) | 0,068 |
| 10 to 15 years | 100 (51,3) | 30 (44,0) | 10 (32,0) | 22 (62,9) | 32 (56,1) |  |
| **Schooling** |  |  |  |  |  |  |
| Children's Ed | 18 (9,2) | 01 (1,5) | 0 (0,0) | 01 (2,8) | 6 (10,5) | 0,232 |
| Elementary school | 102 (52,3) | 40 (58,8) | 25 (80,7) | 17 (48,6) | 26 (45,6) |  |
| Middle school | 75 (38,5) | 27 (39,7) | 06 (19,3) | 17 (48,6) | 25 (43,9) |  |
| **Localization** |  |  |  |  |  |  |
| Urban area | 149 (76,4) | 57 (84,0) | 29 (93,5) | 30 (85,7) | 52 (91,2) | 0,130 |
| Countryside | 46 (23,6) | 11 (16,0) | 2 (6,45) | 05 (14,3) | 05 (8,8) |  |

\* Calculation of schoolchildren with spots made from the total number of schoolchildren with spots with
defining characteristics (n = 329). \*\* Pearson's Chi-square test of schoolchildren with dormant spots.

While schoolchildren who scored on the self-image record had dormant spots, 51.4% (18/35) were male schoolchildren, 62.9% (22/35) were in the age range of 10 to 15 years, and 48, 6% (17/35) attended Elementary School I and the same percentage in Elementary School II, and 85.7% (30/35) resided in the urban zone (Table 2).

The profiles of schoolchildren with suspicious spots for leprosy, 39.2% (129/329) of schoolchildren who have spots, but not from birth, itch and does not aches. Of the schoolchildren with suspicious spots 7.9% (26/329) have cases of leprosy in the family. And schoolchildren with suspicious dormant spots represent 6.7% (22/329) and of these 2.4% (8/329) have cases of leprosy in the family (Table 3).

**Table 3.**
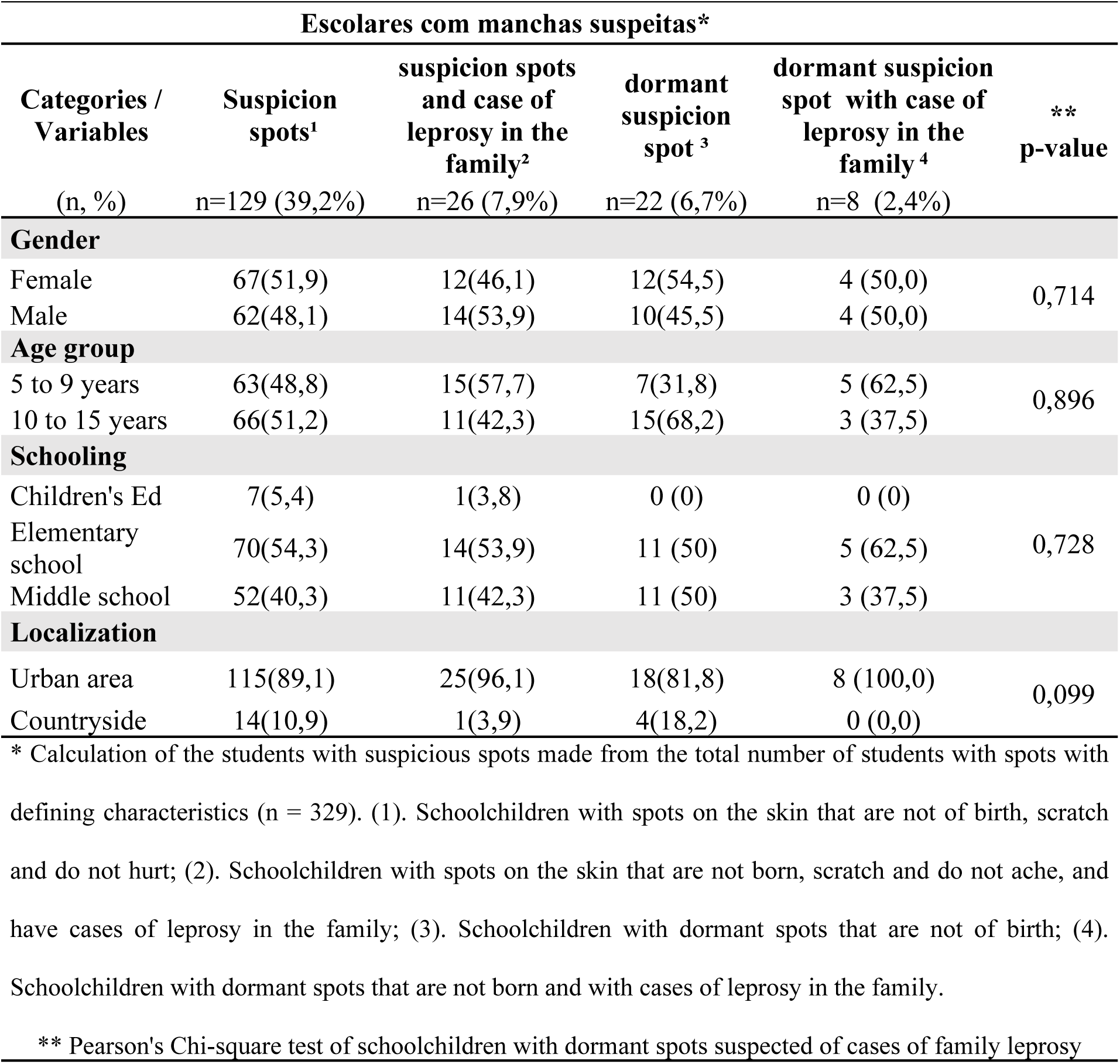
Clinical and epidemiological characteristics of students with suspected leprosy in the IV National Leprosy Campaign in Sobral, Ceará, Brazil, 2018.

The concentration of schoolchildren with suspicious spots was 51.9% (67/129), and 51.2% (66/129) were in the age range of 5 (five) to 9 (nine) years, 54, 3% (70/129) were in Elementary School I and 89.1% (115/129) were residing in the urban area. And schoolchildren with suspicious spots with cases of leprosy in the family, 53.9% (14/26) are males, 57.7% (15/26) are in the age group from 5 (five) to 9 (nine) 96.1% (25/26) lived in the urban area (Table 3).

Among schoolchildren with suspicious dormant spots, 54.5% (12/22) of schoolchildren are female, 68.2% (15/22) were between 10 and 15 years of age, with 50% (11/22).) in Elementary School I, and an equal percentage in Elementary School II, with 81.8% (18/22) residing in the urban area. While students with suspicious dormant spots with cases of leprosy in the family who were screened for the diagnostic examination have the same percentage of participation with 50% (4/8) of male and female schoolchildren, 62.5% (5 / 8) are in the age group of five to nine years and attend Elementary School I, and 100% (8/8) of the schoolchildren lived in the urban zone (Table 3).

### Spatial Analysis of Cases

The evaluation of data from the 24 districts of Sobral, based on the cartographic base of the IBGE, with the mapping of the respective endemicity rates of leprosy in 2016, showed that: 29.2% (7/24) of the neighborhoods have a profile hyperendemic for leprosy; 12.5% (3/24) have a very high rate and have the same high rate percentage; 8.3% (2/24) have an average rate for leprosy; and 37.5% (9/24) did not present any case of leprosy registered in 2016.

In Figure 1, the group of schoolchildren who had spots on the body (34.9%, 317/909) (Map1A), the linear distance matrix among schoolchildren who have cases of leprosy in the family has a minimum distance of 0.0km and the maximum of ± 6.03km; and distance for schoolchildren with spots and cases of leprosy in the family (19.6%, 178/909) (Map 1B) is 0.0km the minimum distance, and the maximum distance is ± 0.041km. The distance between schoolchildren with spots and cases of leprosy in the family (65,1; 592/909) (Map1C) and schoolchildren without spots, the minimum distance is ± 0.0km and the maximum ± 9.1km. The minimum distance between schoolchildren with different characteristics, such as having spots or presenting cases of leprosy in the family, corresponded from 0.0 km to 10 km.

**Figure 1.**
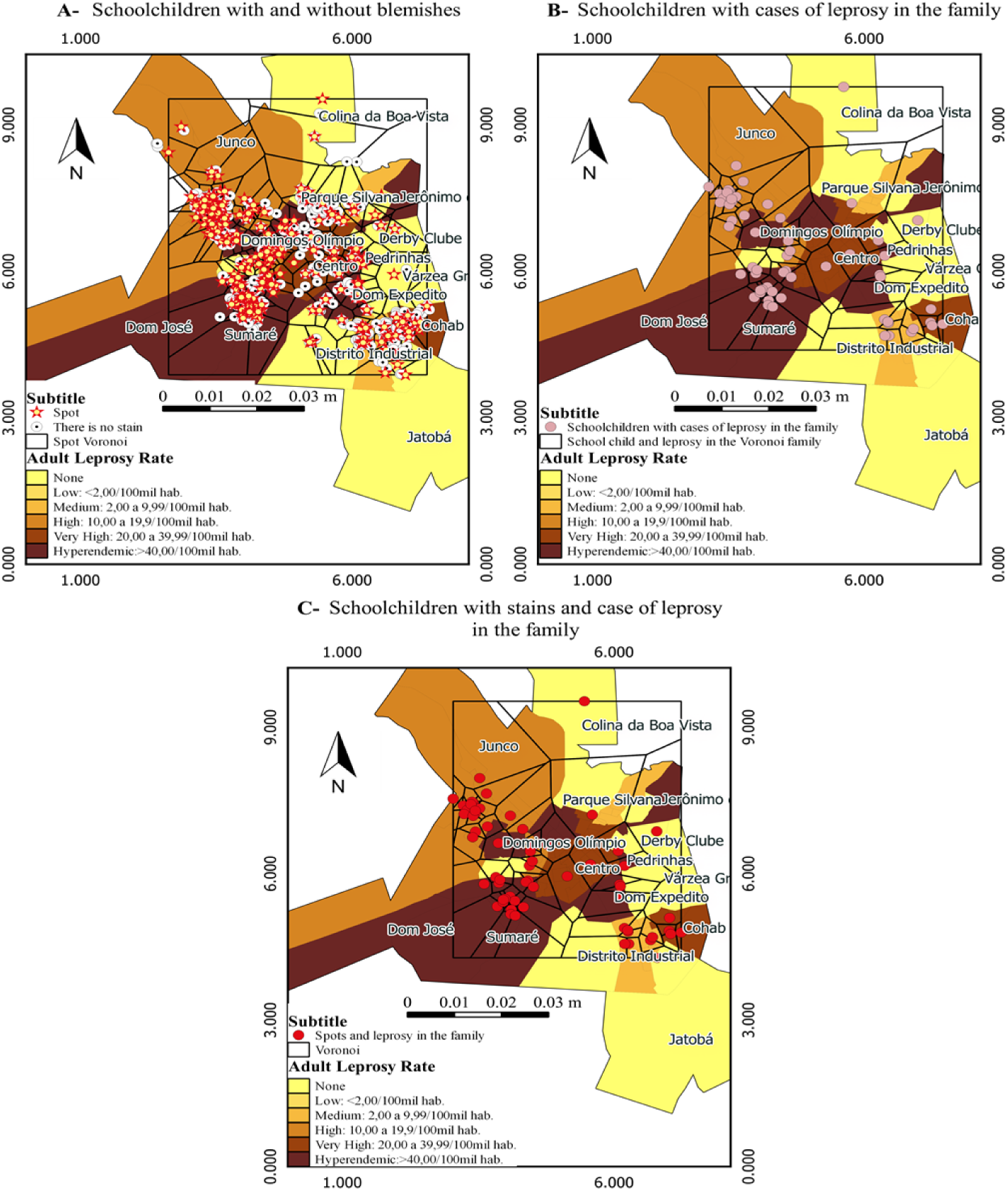
Voronoi maps (Map 1A, 1B and 1C) with the spatial distribution of schoolchildren by neighborhood of residence with their respective leprosy detection rates in 2016, and the characteristics investigated for the presence or absence of spots in the body, and have or do not have cases of leprosy in the family. Source of shapefile: Brazilian Institute of Geography and Statistics (IBGE in Portuguese; https: ftp://geoftp.ibge.gov.br/organizacao_do_territorio/malhas_territoriais/malhas_municipais/municipio_2015/UFs/CE/).

Schoolchildren with spotting who presented cases of leprosy in the family (Map1C), have a neighborhood index of ± 0.6km, and are concentrated the distribution in the neighborhoods that present hyperendemic rates, very high and high leprosy. Only three of the cases with cases of leprosy cases in the family are distributed in three regions where the incidence rate for leprosy was zero, and the respective neighborhoods are: Colina da Boa Vista, Cohab I and Derby.

Figure 2 shows the group of schoolchildren with dormant spots (Map 2 A, 3.8%, 35/909), has a neighbor index closer to ± 0.75km between them, and this group presents a minimum distance of 0.0km and a maximum of ± 0.06km in relation to schoolchildren suspected of leprosy (20%, 7/35) (Map2B).

**Figure 2.**
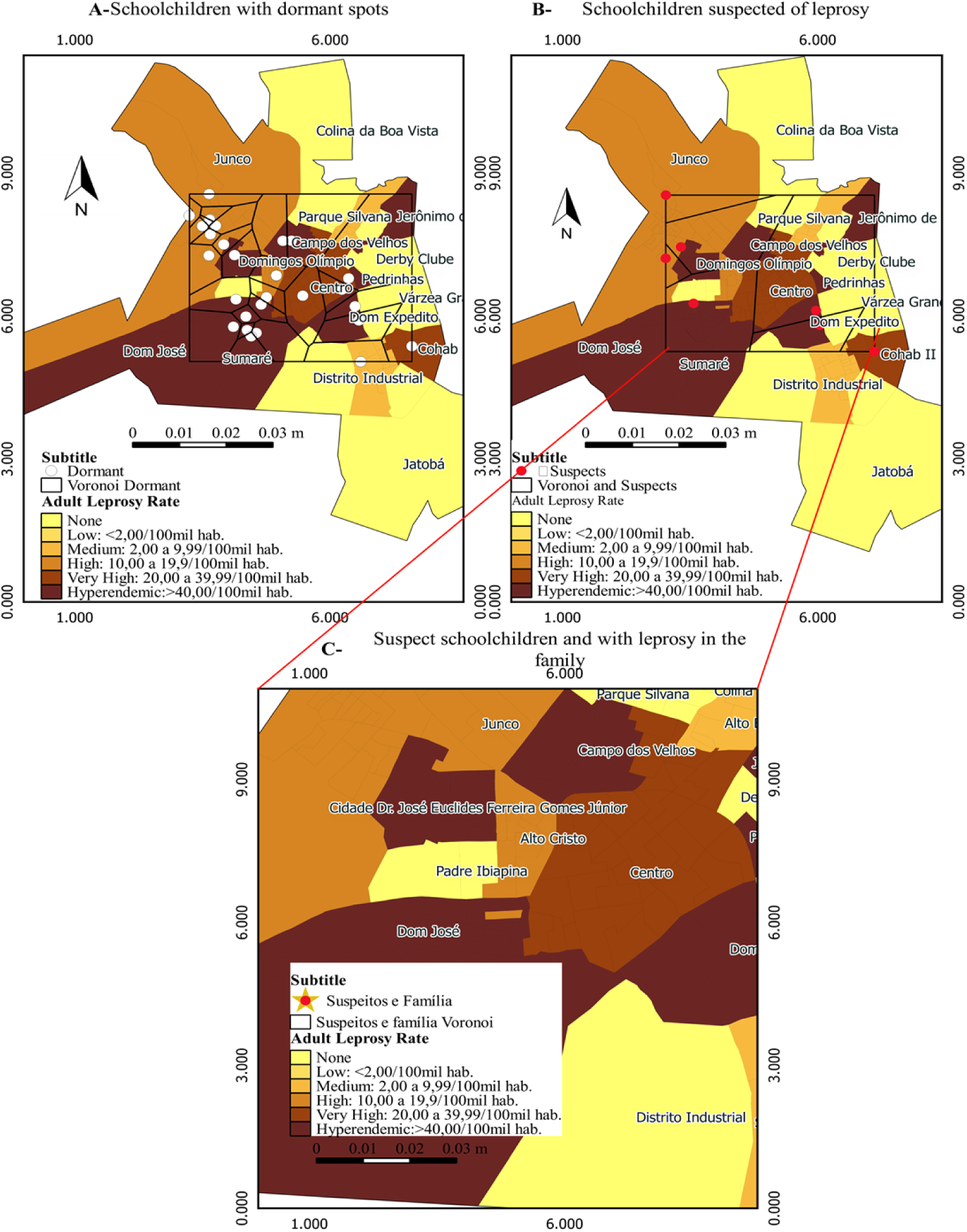
Voronoi maps (2A, 2B and 2C) with the spatial distribution of leprosy detection rates in Sobral in the year 2016, of schoolchildren with dormant spots suspected of leprosy and cases of the disease in the family. Source of shapefile: Brazilian Institute of Geography and Statistics (IBGE in Portuguese; https: ftp://geoftp.ibge.gov.br/organizacao_do_territorio/malhas_territoriais/malhas_municipais/municipio_2015/UFs/CE/).

The group of schoolchildren suspected of leprosy (Map 2B) has a neighborhood index of ± 0.99km, with an average of three schoolchildren. Of the seven suspected schoolchildren, all are located in neighborhoods with a high leprosy rate for leprosy, with two schoolchildren in the Cidade Dr. José Euclides Ferreira Gomes neighborhood, two schoolchildren at Dom Expedito, and one in Junco and one schoolchildren in the. neighborhood with very high rate for leprosy, which is the Cohab II and a schoolchildren in the hyperendemic neighborhood, the Dom José.

Of the schoolchildren suspected of having leprosy cases in the family (Map C 2, 42.8, 3/7), they have a neighbor index of ± 2.66km, of which three are suspected, two are located in neighborhoods with high leprosy, and a schoolchildren in a hyperendemic neighborhood, being respectively two in the Cidade Dr. José Euclides Ferreira Gomes neighborhood and one in the neighborhood of Dom José.

## DISCUSSION

The National Leprosy Campaign is a successful strategy worldwide referenced by the WHO to act in the active search for new cases of leprosy with schoolchildren in the age group of five to fourteen years old of the public education network residing in areas at highest risk for the disease^15^. The active search strategy consists of epidemiological investigation actions, such as campaigns and surveys^16^.

In Sobral, the National Leprosy Campaign in 2016 had a predominance of female schoolchildren, a result similar to the active search for cases of leprosy in the city of Cuiabá, in the state of Mato Grosso, developed in a elementary school with 10 to 14 year old schoolchildren, where the female participation was 58.6% in a group of 1,263 students^17^.

When analyzed the predominance of the schoolchildren participating in the campaign with spots on the body, the highest concentration was of male schoolchildren. Although the presence of the lesion does not confirm whether it is a case of leprosy, since it has to meet the cardinal signs of a case definition, studies show that men are more affected by leprosy than women^18^. In Brazil, in na temporal analysis study distribution of leprosy by sex in the period 2012-2016, presented similar results from the WHO, even from the age of 15 years old.

Leprosy elimination guidelines define leprosy cases when one of the three cardinal signs is verified: 1 – Lesions with altered sensitivity, thermal and painful and tactile; 2 – Thickening of nerves, with alteration; 3 – Presence of M. leprae confirmed in skin crevices ^15, 20^.

In the analysis of the sample, it was also possible to investigate some epidemiological aspects of the t schoolchildren in relation to cases of leprosy in the family nucleus. When children are exposed to M. leprae in the family environment, they have a 60% higher chance of developing leprosy. If the child has contact with cases of leprosy in the environment outside the home is four times more likely to develop leprosy, although the development of the disease depends on the relationship of the bacillus and host ^15, 21, 22^. In an active search campaign of new leprosy cases in Manaus (2018) were detected 40 new cases of leprosy in schoolchildren, 21 of whom had household contacts of leprosy transmission sources, and 95.2% had family contact with someone who had or has leprosy ^23^.

As a strategy to enhance early detection and tracing of new cases of leprosy, spatial analysis has its place in epidemiology as an important tool for understanding the dynamics of leprosy transmission, by directing the active search for new cases ^24, 25^. Thus, the Geographic Information System (SIG) provides a spatial analysis of the distribution of the characteristics of the spots of the schoolchildren with the rate of detection of leprosy by census tracts, focusing on schoolchildren with dormant spots suggestive of leprosy.

The maps allow analysis and understanding of spaces, social relations and landscape ^26^. The construction of the maps allowed to identify and analyze from the characteristics of the records of self-image correlations with endemic areas for leprosy in the neighborhoods of Sobral.

From the distribution of the endemicity rate of new leprosy cases registered in 2016, it was verified that 29.2% of Sobral’s neighborhoods have a hyperendemic profile for leprosy, which makes the resident population with the potential to develop M. leprae, and reinforces the need for the development of strategic actions of active search, combat and control, since the number of cases registered in passive search may be greater when performing active search actions ^27^.

Brazil has a heterogenic spatial distribution of cases of leprosy, guided by the economic social level. Southern states with greater socioeconomic development do not reach high levels or hyperendemicity of leprosy. Meanwhile, states of the North, Midwest and Northeast still face high endemic loads ^28, 29^.

In the analysis with the Voronoi Maps, the study identified that schoolchildren who have spots on the body maintain distances close to those who have cases of leprosy in the family, reaching a minimum of zero kilometer. The study carried out in the State of Mato Grosso found that 12 new cases diagnosed from a set of 34 cases between 2001 and 2007 have a distance of 50 meters from another case diagnosed with leprosy ^30^.

The physical distance and the genetics of confirmed leprosy cases are considered risk factors for the development of leprosy in healthy people. These factors should be considered in the social relations between susceptible and vulnerable people within the neighborhood, due to the risk of developing leprosy in healthy people from a family contact is nine times as much ^31, 32, 28^.

The dissemination of information that characterizes leprosy may prompt the population to seek diagnosis earlier, such as facilitating active search campaigns. Leprosy has a high capacity to cause permanent and irreversible lesions and the detection of the disease in children under 15 years old is an indicator of sensitivity for transmission of M. leprae, implying maintenance of the bacillus transmission cycle and failures in monitoring and control policies ^33, 34^.

## CONCLUSION

The active search strategy adopted by the National Leprosy Campaign is an effective methodology for screening suspected cases among schoolchildren and early detection of new cases of leprosy, preventing physical disabilities resulting from late diagnosis. Early detection of leprosy and immediate treatment to avoid disability minimizes stigma associated with leprosy such as discrimination, as many patients continue to suffer from social exclusion resulting from leprosy injury.

It should be noted that, in the Sobral Leprosy Campaign, no new cases of leprosy were found in children under 15 years of age in the campaign, despite the schoolchildren of the students with dormant and suspicious spots, a profile of suspected schoolchildren with cases of leprosy in the family residing in areas with high rates of leprosy. However, the campaign allowed to establish the epidemiological characteristics of the participating students, such as stratification of groups residing in areas with high endemic rates.

This study had several limitations, first because it is a study with a secondary data source, being only possible the access of the self-image files returned by the students to the school officials during the period of the National Leprosy Campaign. Second, in addition to the losses of return of the self-image files for leprosy, an expressive quantity of fichas was not correctly filled, being excluded from the study and reducing the sample for analysis. Thus, there is a need to raise the awareness of those responsible for the importance of returning the self-image record, in order to evaluate new cases suspected of leprosy in <15 years.

